# Functional modulation of T follicular cells *in vivo* enhances antigen-specific humoral immunity

**DOI:** 10.1101/2020.11.17.387100

**Authors:** Jose D. Pagan, Hera Vlamakis, Anthony Gaca, Ramnik Xavier, Robert M. Anthony

## Abstract

Generation of high-affinity IgG is essential for defense against infections and cancer, is the intended consequence of many vaccines, but can cause autoimmune and inflammatory diseases when inappropriately directed against self (Wang et al., 2018, Ludwig et al., 2017, Chinen et al., 2010). The interplay and balance of T follicular helper cells (T_FH_) and T follicular regulatory cells (T_FR_) is critical for production of high-affinity IgG (Wing et al., 2018). Here, we empowered T_FH_ cells and improve antigen-specific IgG responses with two interventions intended to transiently diminish T_FR_ influence. First, adult mice were administered an antibiotic cocktail (ABX) for an extended period to deplete the immunoregulatory intestinal microbiota (Belkaid and Harrison, 2017, Thaiss et al., 2016, Rooks and Garrett, 2016, Honda and Littman, 2016, Perruzza et al., 2017, Teng et al., 2016, Block et al., 2016, Proietti et al., 2014, Slack et al., 2014). This treatment skewed T follicular cell ratios, with increased T_FH_ and reduced T_FR_ numbers. TNP-KLH immunization resulted in higher affinity TNP-specific IgG in ABX mice compared to controls. In a model of IgG-driven inflammatory nephritis, ABX mice had significantly worse nephritis accompanied by higher affinity antigen-specific IgG, and enriched T_FH_ cells compared to controls. Second, we sought to functionally manipulate T_FH_ and T_FH_ cells, which both express the checkpoint inhibitory molecule, PD-1 (Sage et al., 2013), by administration of α-PD-1 during immunization. This intervention enhanced the affinity of antigen-specific IgG and increased in T_FH_ following TNP-KLH immunization and nephritis induction. These results suggest that altering T_FH_ and T_FR_ ratio during immunization is an appealing strategy to qualitatively improve IgG responses.

## Introduction

Immunoglobulins (IgG) link the adaptive and innate immune systems with their two domains that combine precise specificity for antigen in the antigen-binding fragment (Fab), and convey effector functions via the constant, crystallizable domain (Fc) interactions with Fc γ receptors and complement (Schroeder and Cavacini, 2010, Nimmerjahn and Ravetch, 2008, Ravetch and Bolland, 2001). Production of high-affinity antigen-specific IgG of appropriate subclass is essential for host defense against many infections, and the intended consequence of many vaccines (Zimmermann and Curtis, 2019, Compeer and Uhl, 2020, Robbiani et al., 2020, Yang et al., 2020).

Insights into the cellular and molecular regulation of IgG production have highlighted the role of follicular-resident T cells as crucial gate keepers (Wu et al., 2016). T follicular helper cells (T_FH_) provide B cell help to drive IgG production, whereas T follicular regulatory cells (T_FR_) set activation thresholds for these responses (Crotty, 2011, Hatzi et al., 2015, Wu et al., 2016). T_FH_ cells are identified in by surface expression of CD4, PD1, and CXCR5, and the transcription factor expression, Bcl6 (Hatzi et al., 2015). These markers are shared with T_FR_, which also express Foxp3 (Sage et al., 2013). Thus, striking a balance between these cells is critical for productive, and not harmful, IgG responses. Indeed, some autoimmune diseases results from IgG generated against host antigens, underscoring the importance of the balance between T_FH_ and T_FR_ cells (Zhang et al., 2017, Sage and Sharpe, 2015, Wang et al., 2020).

The intestinal microbiota plays critical roles in development and regulation of the immune system (Rooks and Garrett, 2016). Germ-free mice lacking any bacteria have severely restricted T cell and B cell repertoires (Chen et al., 2018, Wesemann et al., 2013, Li et al., 2020). Further, the microbiota influences the immune system and its responses through multiple non-mutually exclusive mechanisms. Microbiota-derived metabolites, including short-chain fatty acids (SFCA), have been shown to promote T regulatory cells (Chinen et al., 2010, Atarashi et al., 2011, Marino et al., 2017) and also T_FR_ cells (Takahashi et al., 2020). Furthermore, depletion of the microbiota revealed exacerbated or harmful immune responses, generally attributed to T regulatory deficiencies or deficits (Atarashi et al., 2011, Belkaid and Harrison, 2017, Chinen et al., 2010, Honda and Littman, 2016, Marino et al., 2017, Maslowski et al., 2009, Rakoff-Nahoum et al., 2004, Rooks and Garrett, 2016).

In contrast to this, depletion of the microbiota in humans by a 5-day treatment with broad-spectrum antibiotics led to impaired IgG responses to flu vaccination (Hagan et al., 2019). Because the effects of dietary SFCA on the immune system required weeks of feeding (Marino et al., 2017, Maslowski et al., 2009, Tan et al., 2017), we explored the effect of an extended antibiotic treatment on immunization in addition to characterizing antibody subclass response. We therefore asked whether treatment with broad-spectrum antibiotics for an extended period would impact T_FH_ and T_FR_ cells, service as an initial platform to influence IgG responses.

## Results

We set out to characterize the effect of long-term microbiota depletion on immune response, and orally administered a cocktail of ampicillin, neomycin, metronidazole, and vancomycin (Rakoff-Nahoum et al., 2004), collectively referred to as ABX for several weeks. Fecal samples from treated animals were cultured under aerobic conditions on Luria broth (LB) agar and anaerobic conditions on brain heart infusion (BHI) agar to reveal a significant reduction in colonies in ABX-mice compared to water-treated wild type (WT) control animals (Figure 1A). 16S sequencing of fecal pellets confirmed marked perturbations of the microbiota by ABX, as the dominance of *Bacteroidetes* and *Firmicutes* in WT was replaced by *Firmicutes* and *Proteobacteria* (Figure 1B, Figures S1A-B). Serum IgG antibody titers in ABX and WT mice were assessed in ELISA, revealing similar total mouse IgG1 (mIgG1, Figure 1C), increased total mIgG2a/c (Figure 1D), increased total mIgG2b (Figure 1E), decreased total mIgG3 (Figure 1F), increased total mIgE (Figure 1G), similar total mIgM (Figure 1H), decreased total mIgA (Figure 1I), as well as total fecal mIgA (Figure 1J) in ABX-mice compared to WT controls. Intriguingly, the extended ABX treatment shifted homeostatic total mIgG production towards more inflammatory subclasses, including mIgG2a/c and mIgG2b.

**Figure 1.**
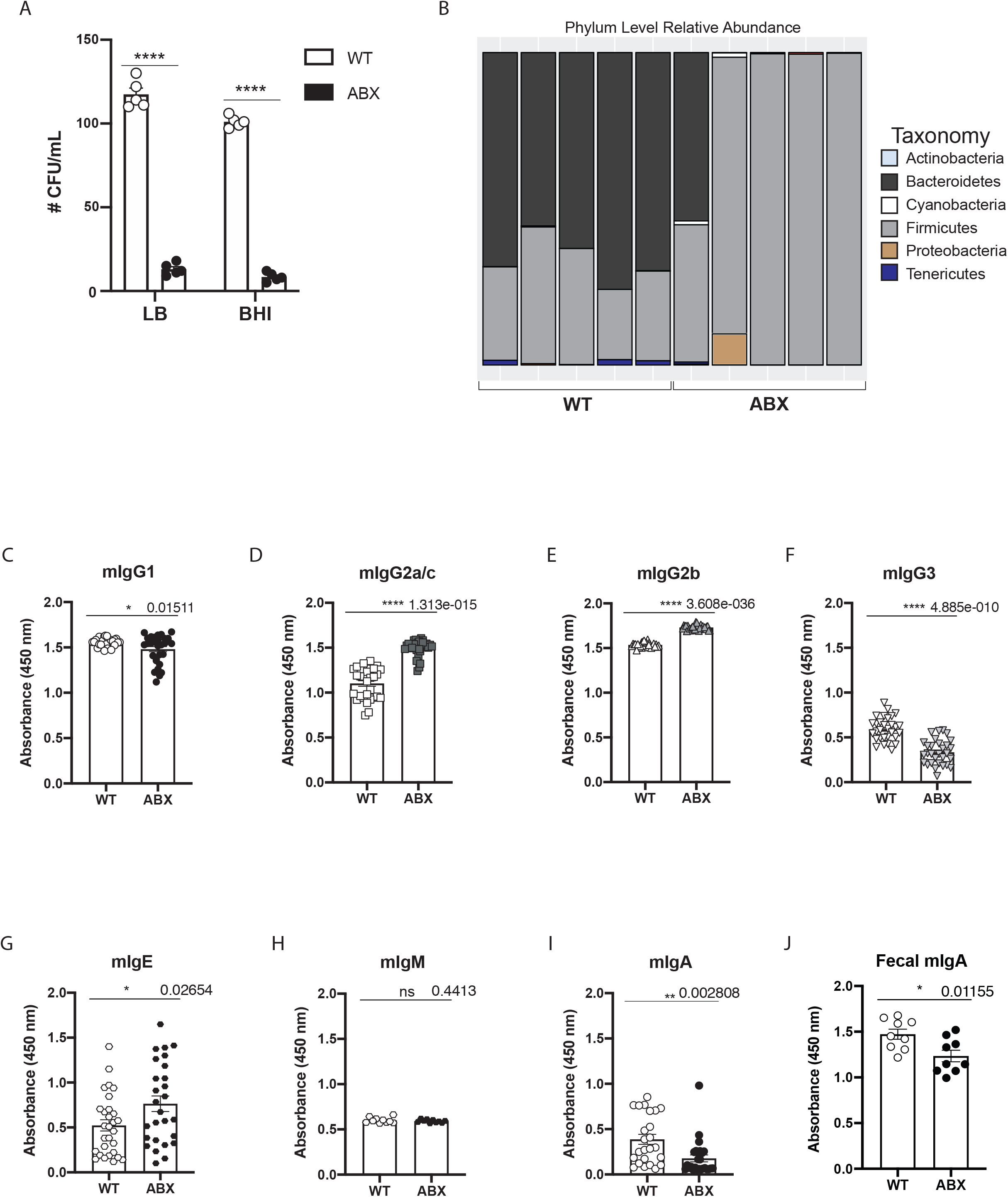
ABX-treatment alters IgG subclass distribution during homeostasis. (**A**) Colony forming units (CFU) grown on LB agar and brain heart infusion (BHI) agar plates from fecal bacteria after antibiotic treatment of mice (*n* = 5 mice/group). (**B**) Stool samples sequenced using 16S data and analyzed to determine phylum level relative abundance. Serum immunoglobulin titers of WT (white filled) and ABX treated mice (black filled) for (**C**) mIgG1 (*n* = 20-30), (**D**) mIgG2a/c (*n* = 20-30), (**E**) mIgG2b (*n* = 20-30), (**F**) mIgG3 (*n* = 20-30), (**G**) mIgE (*n* = 20-30), (**H**) mIgM (*n* = 10), (**I**) mIgA (*n* = 20), and (**J**) fecal mIgA (*n* = 10) determined by ELISA. Means and standard error of the mean (±SEM) are plotted. Results are representative of at least two independent repeats. *p<0.05, **p<0.01, ***p<0.005, ****p<0.001, ns (not significant), p values are for Two-tailed unpaired *Student’s* t test were performed for statistical comparisons.

We asked whether ABX altered lymphocyte populations, leading to altered homeostatic total antibody titers (Belkaid and Harrison, 2017, Chen et al., 2018, Chinen et al., 2010, Wesemann et al., 2013). Splenic lymphocyte populations involved in IgG production were examined in WT control and ABX-mice by flow cytometry (Figure S2). No difference between these groups was apparent in CD19^+^ B cells (Figure 2A-C), CD3^+^ T cells (Figure 2A-C), follicular B cells (FoB; CD23^+^ CD21/35^+hi^ IgM^lo^; Figure 2D-F, Figure S1A-D), transitional 2 B cells (T2; CD32^+^ CD21/35^+^ IgM^+^; Figure 2D-F, Figure S2A), marginal zone B cells (MZB; CD19^+^ CD23^−^ CD21^+^ CD35^med^; Figure 2G-I)) transition 1 B cells (T1; CD19^+^ CD23^−^ CD21^−^ CD35^med^; Figure 2G-I). However, germinal center B cells (GC, B220^+^ PNA^+^ FAS^+^, Figure 2J and K, Figure S2B) and T follicular helper cells (T_FH_, CD4^+^ CXCR5^+^ PD1^+^, Figure 2L and M, Figure S2C-D) were enriched in ABX-treated mice compared to WT controls, while T follicular regulatory cells (T_FR_, CD4^+^ FoxP3^+^ CXCR5^+^ PD1^+^, Figure 2L and N, Figure S2C-D) were reduced, resulting in an altered T_FR_:T_FH_ ratio (Figure 2O).

**Figure 2.**
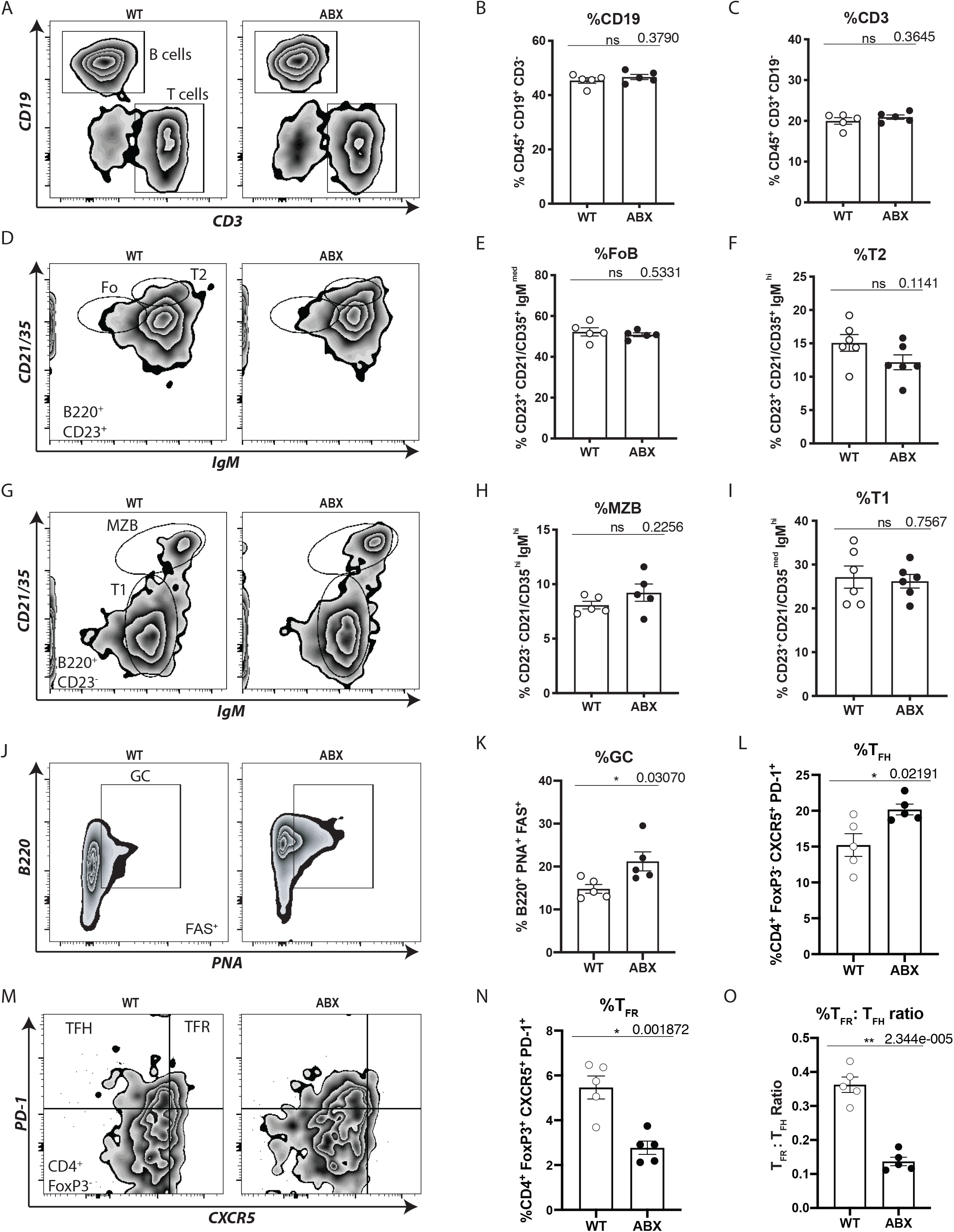
ABX enriches Germinal Center B cells and T follicular help cells. Representative flow cytometric analysis of splenic **(A)** B cells (CD19^+^) and T cells (CD3^+^), and quantification of CD19^+^ cells (**B**), CD3^+^ T cells (**C**). Representative plots for follicular B cells (CD21/35^+^ IgM^Med^) gated on B220^+^ CD23^+^ cells, and transitional type 2 B cells (CD21/35^+^ IgM^High^) gated on B220^+^ CD23^+^ cells (**D**), and quantification follicular B cells (**E**) and Transitional type 2 B cells (**F**). **(G)** Representative plots of Transitional type 1 B cells (CD21/35^Low^ IgM^Med^) gated on B220^+^ CD23^−^ cells and marginal zone B cells (CD21/35^High^ IgM^High^) gated on B220^+^ CD23^−^ cells, and quantification of marginal zone B cells (**H**) and Transitional type 1 B cells (**I**). **(J)**Representative Germinal Center B cells (GC, FAS^+^ PNA^+^) gated on B220^+^ cells and their frequency (**K**). **(L)** Representative FACS plots of T_FH_ cells (FoxP3^−^ PD-1^+^ CXCR5^+^) and T_FR_ (FoxP3^+^ PD-1^+^ CXCR5^+^) gated on CD4^+^ cells, and their quantification (**M, N**). Means and standard error of the mean (±SEM) are plotted. Results are representative of at least two independent repeats. *p<0.05, **p<0.005, ****p<<0.001, ns (not significant), determined by two-tailed unpaired *Student’s* t test (*n* = 5 mice/group).

Because T_FH_ cells and GC B cells were enriched during extended ABX-treatment, we turned to a model antigen immunization system to characterize antigen-specific IgG responses. ABX- and WT control mice were immunized with 2,4,6-trinitrophenyl hapten conjugated to keyhole limpet hemocyanin (TNP-KLH) and alum, and the total and TNP-specific IgG response was examined (Figure 3A). On day 7, total mIgG1 titers were reduced in ABX-mice (Figure 3B), mIgG2a/c were unchanged (Figure 3C), and mIgG2b (Figure 3D), and mIgG3 (Figure 3E) were increased. However, TNP-specific IgG titers were similar across all subclasses in WT controls and ABX-animals (α-TNP mIgG1, Figure 3F; α-TNP mIgG2a/c, Figure 3G; α-TNP mIgG2b, Figure 3H; α-TNP mIgG3, Figure 3I). ABX-treatment resulted in an increased percentage of GC B cells (Figure 3J), T_FH_ cells (Figure 3K), decreased T_FR_ cells (Figure 3L), and a decreased T_FR_:T_FH_ ratio (Figure 3M). The affinity of IgG for TNP was examined, revealing ABX-mice produced two-fold higher affinity IgG compared to WT controls (Figure 3N) (Hajighasemi et al., 2004). Taken together, extended ABX treatment increases T_FH_ cells and GC B cells, and results in higher affinity IgG following immunization.

**Figure 3.**
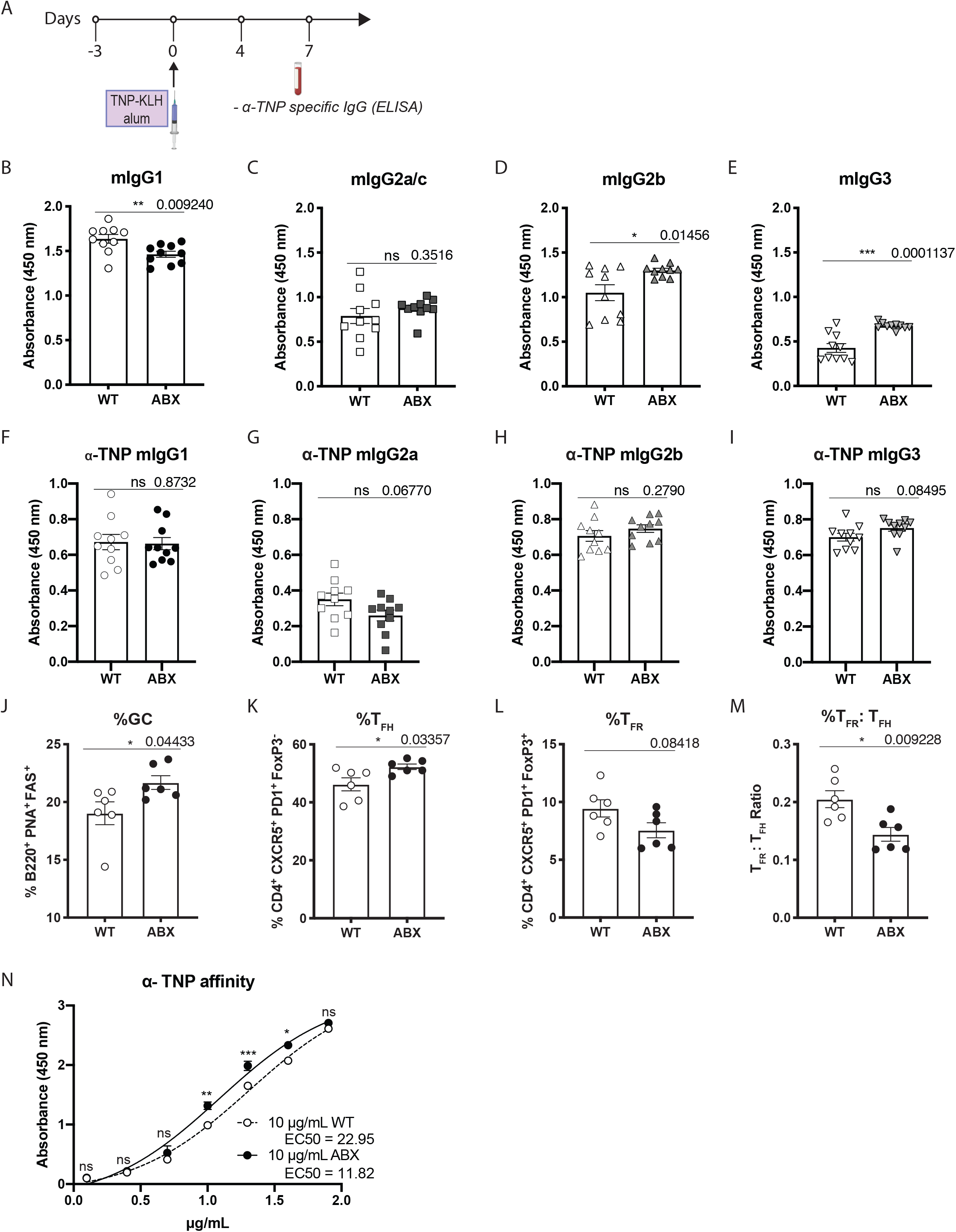
ABX enhances TNP-specific responses. (**A**) Schematic of TNP-KLH immunization in alum. Quantification of subclass-specific total IgG after NTN-induction by ELISA for mIgG1 (**B**), mIgG2a/c (**C**), mIgG2b (**D**), and mIgG3 (**E**) (*n* = 10 mice/group). TNP-specific IgG titers on day 7 for mIgG1 (**F**), mIgG2a/c (**G**), mIgG2b (**H**), and mIgG3 (**I**) (*n* = 10 mice/group). Frequency of splenocytes 7 days following TNP-KLH immunization, including germinal center B (**J**), T_FH_ cells (**K**), and T_FR_ cells (**L**). **(M)** Ratio of T_FH_: T_FR_ from **L** and **K** (*n* = 6 mice per group). **(N)** Affinity constant plot of isotype-treated (white circles) treated and ABX-treated (black circles) mice after TNP-KLH immunization (*n* = 3-5 mice/group). Means and standard error of the mean (±SEM) are plotted. Results are representative of at least two independent repeats. *p<0.05, **p<0.005, ****p<0.001, ns (not significant), determined by Student’s t test.

We next explored the functional contribution of extended ABX treatment by a model of induced Goodpasture’s Disease that culminates in IgG-driven acute nephrotoxic nephritis (NTN). WT or ABX-mice were immunized with sheep IgG and Complete Freund’s Adjuvant (CFA; Figure 4A). Four days later, the mice were administered anti-mouse glomeruli basement membrane sera (GBM) raised in sheep (sheep α-GBM), which effectively targets immune complexes to the kidney (Salant and Cybulsky, 1988, Kaneko et al., 2006a). On day 7 following sheep α-GBM treatment, serum was collected and assessed, revealing little differences in total IgG across all subclasses in the sera (total mIgG1, Figure 4B; total mIgG2a, Figure 4C; total mIgG2b, Figure 4D; total mIgG3, Figure 4E). However, analysis of antigen-specific α-sheep mIgG titers in sera showed reduced titers of α-sheep mIgG1 and α-sheep mIgG2b in ABX-treated mice compared to WT controls (Figure 4F-I). Despite this, ABX-mice suffered higher mortality (Figure 4J) and significantly more kidney damage as measured by blood urea nitrogen (BUN) on day 7 (Figure 4K) compared to WT controls, indicating microbiome depletion exacerbated IgG-mediated autoimmune nephritis. These results of reduced titers were consistent to a study of responses to flu vaccination during 5-day ABX (Hagan et al., 2019), but indicated the IgG generated during extended ABX-treatment had enhanced effector function/functionality.

**Figure 4.**
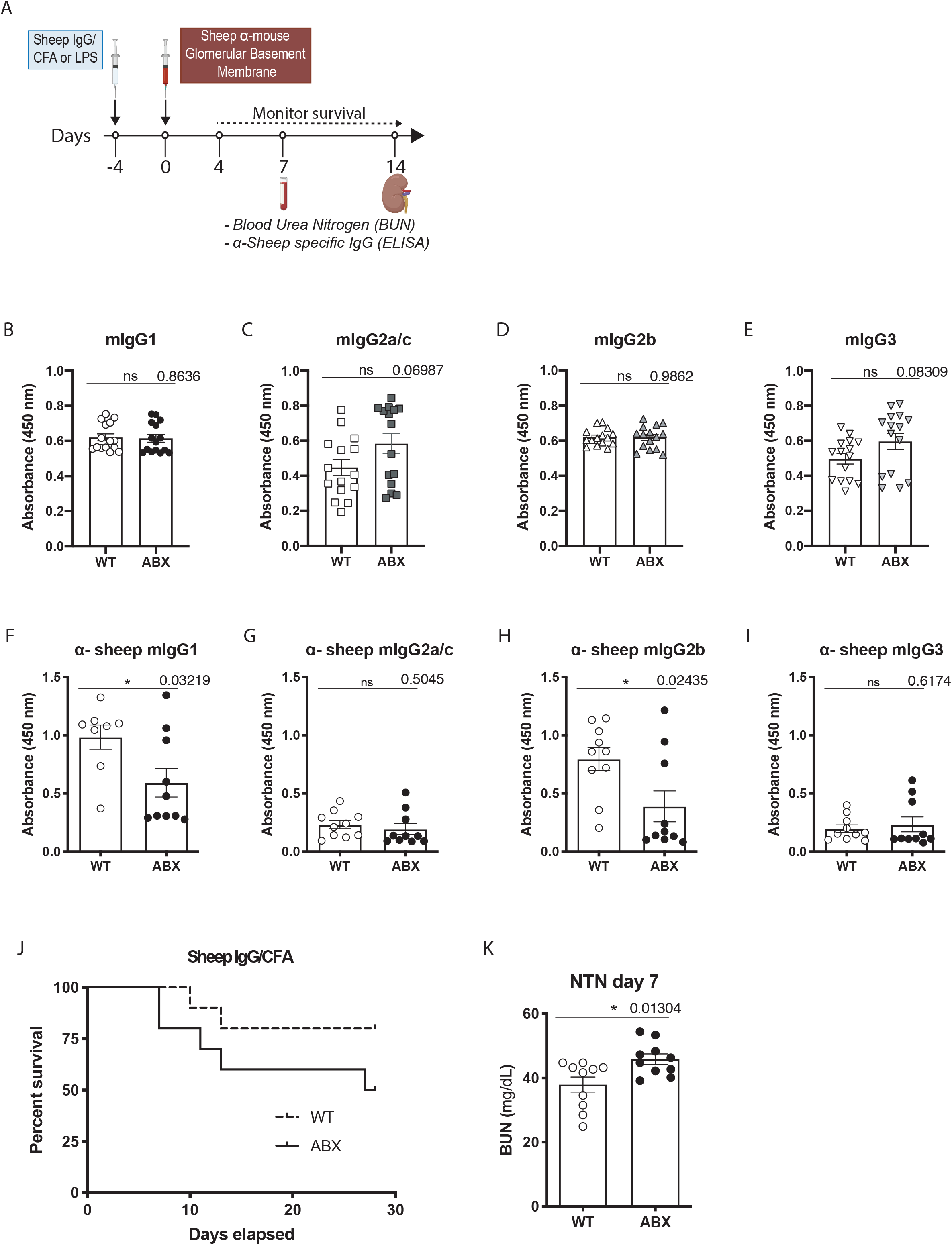
ABX alters IgG titers and disease during induced nephritis. (**A**) Schematic of NTN-induction protocol. Quantification of subclass-specific total IgG after NTN-induction by ELISA for mIgG1 (**B**), mIgG2a/c (**C**), mIgG2b (**D**), and mIgG3 (**E**) (*n* = 10 mice/group). Sheep IgG-specific IgG day 7 after NTN-induction for mIgG1 (**F**), mIgG2a/c (**G**), mIgG2b (**H**), and mIgG3 (**I**) in mice were treated with PBS (white filled), or ABX (black filled) (*n* = 10 mice/group). (**J**) NTN-induced survival curve (*n* = 10 mice/group) and day 7 blood urea nitrogen (BUN) levels on day 7 (*n* = 5 mice/group) (**K**) in PBS-treated and ABX-treated mice. Means and standard error of the mean (±SEM) are plotted. Results are representative of at least two independent repeats. *p<0.05, **p<0.01, ***p<0.005, ****p<0.001, ns (not significant), p values are for unpaired *Student’s* t test.

We next asked whether extended ABX treatment altered effector cells that mediate IgG-responses, including neutrophils and macrophages, thereby compensating for lower IgG titers. Therefore, we passively transferred arthritogenic K/BxN sera (Korganow et al., 1999) to ABX and WT control mice, to induce IgG-mediated joint inflammation. Footpad swelling was monitored over the next several days revealing the transferred sera induced robust arthritic swelling of paws in both treatment groups, although swelling was slightly reduced in ABX animals (Figure S3A) suggesting WT and ABX animals have similar abilities to elicit IgG functions.

We next examined qualitative differences in mIgG produced by ABX animals during NTN using LPS instead of CFA, because extended ABX treatment enhanced affinity of antigen-specific IgG (Fig. 3N). Analysis of sera mIgG from NTN-induced WT controls and ABX mice revealed similar steady-state binding affinities for sheep IgG (Figure 5A). Because NTN targets immune complexes to the kidney, we compared mIgG from the kidney of NTN-induced WT controls and ABX-treated mice. Intriguingly, mIgG recovered from the kidney of ABX mice had a marked higher affinity for sheep IgG than kidney-derived mIgG from WT controls (Figure 5B). This trend held across mIgG subclasses, as kidney α-sheep mIgG1 (Figure 5C), α-sheep mIgG2a/c (Figure 5D), and α-sheep mIgG2b (Figure 5E) from NTN-induced ABX mice had higher affinity for sheep IgG compared to WT controls.

**Figure 5.**
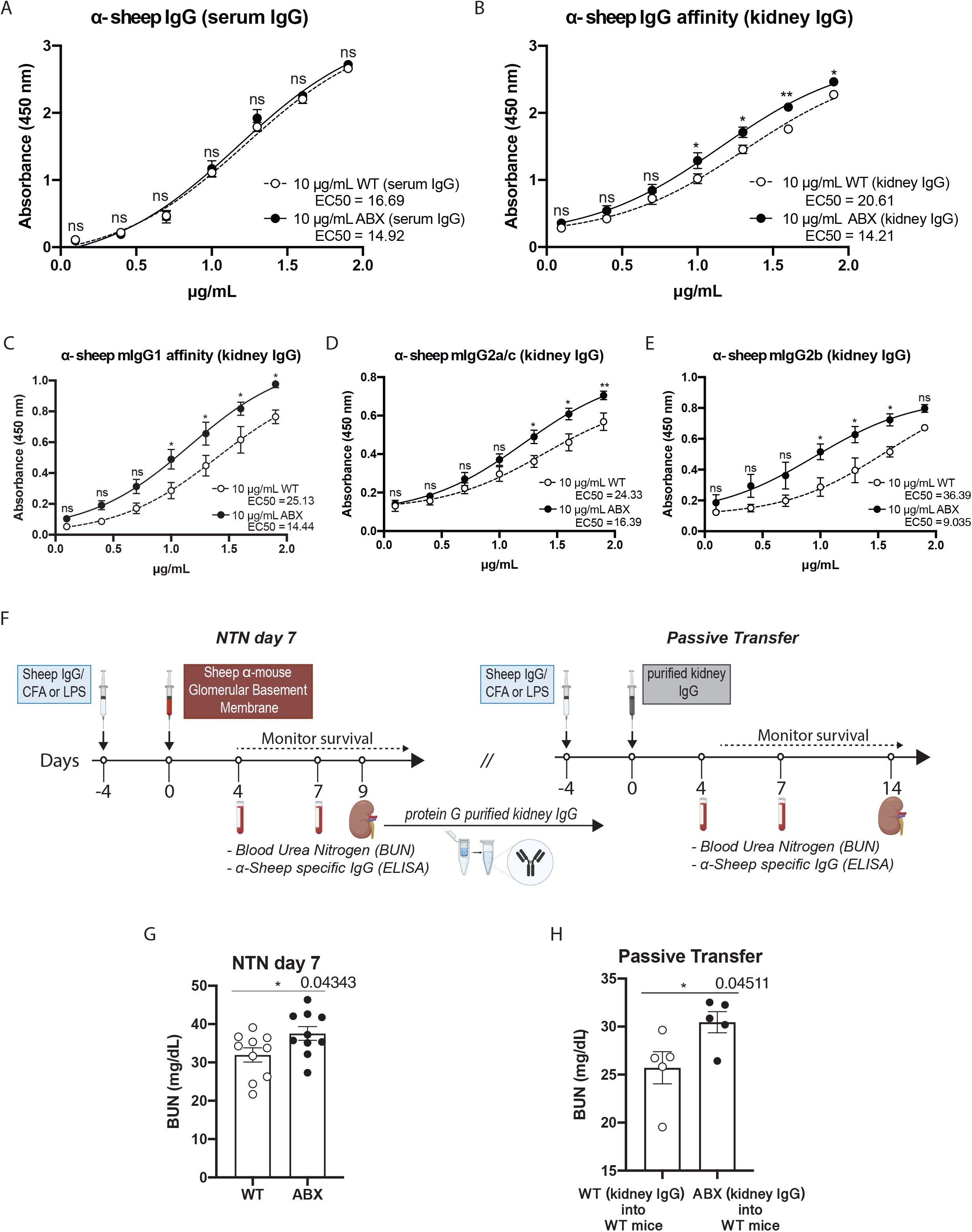
Enhanced potency of kidney-deposited IgG in ABX mice. (**A, B**) Sheep IgG-specific affinity constant plot sera-derived IgG from PBS-treated (white circles) and ABX-treated (black circles) mice after NTN-induction. (**A**) α-sheep IgG affinity plot of total IgG antibodies recovered from PBS-treated (white filled) and ABX-treated (black filled) mouse serum on day 9 following NTN-induction (*n* = 5 mice/group). (**B**) α-sheep IgG affinity plot of total IgG antibodies from kidneys of PBS-treated (white filled) and ABX-treated (black filled) mice recovered at day 9. Affinity plots of day 9 kidney-derived α-sheep IgG1 (**C**), α-sheep IgG2a/c (**D**), and α-sheep IgG2b (**E**) (*n* = 5 mice/group). (**F**) Schematic for kidney-derived NTN-passive transfer model. (**G**) Blood urea nitrogen (BUN) levels on day 7 of NTN-induction of PBS-treated (white filled) and ABX-treated donor mice (black filled) (*n* = 10 mice/group). **(H)**Blood urea nitrogen (BUN) levels on day 7 in recipient mice following passive transfer of kidney-derived IgG from PBS-treated (white filled and ABX-treated (black filled) donors (*n* = 5 mice/group). Means and standard error of the mean (±SEM) are plotted. Results are representative of at least two independent repeats. *p<0.05, **p<0.01, ***p<0.005, ****p<0.001, ns (not significant), p values are for two-tailed unpaired *Student’s* t test.

To determine whether higher affinity IgG deposited in the kidney of ABX-treated mice following NTN induction was responsible for severity of disease, we passively transferred kidney-deposited IgG (Figure 5F, Figure S4). NTN was induced in WT controls and ABX-treated mice, and confirmed as day 7 BUN levels enhanced in ABX-treated and reduced α-sheep IgG2b responses compared to controls, and similar α-sheep IgG affinity in sera (Figure 5G, Figure S4A-F). IgG was purified from the kidneys of WT and ABX mice at day 9 after α-GBM treatment, and transferred to WT control recipients (Figure 5F, Figure S4E-F). Consistently, WT mice receiving ABX-treated kidney IgG had elevated BUN levels compared to mice administered WT kidney IgG (Figure 5H), despite transfer of equivalent amounts of IgG. These results confirm IgG deposited in the kidney of ABX-animals after NTN-induction is more pathogenic.

Although extended ABX promoted increased GC Bells, increased T_FH_ cells, and higher affinity antigen-specific IgG in two independent immunization models, it is a highly impractical immunization strategy. Exploring alternate strategies, we reasoned that blockade of the inhibitor receptor PD-1 (Agata et al., 1996, Freeman et al., 2000, Ishida et al., 1992) might recapitulate this phenotype. Both T_FH_ and T_FR_ cells express PD-1 (Agata et al., 1996, Ishida et al., 1992, Sage et al., 2013). Indeed, PD-1 blockade is established to license CD8^+^ T cell function in a number of cancers, and α-PD-1 blocking IgG are readily available in the clinic (Iwai et al., 2005, Wei et al., 2019, Wei et al., 2017). Further, while blocking PD-1 promotes CD8+ T cell-driven cytotoxic responses (Freeman et al., 2000, Gubin et al., 2014, Mamalis et al., 2014, Wei et al., 2017), its effect on the humoral response is not fully understood. Mice were administered an abbreviated treatment schedule of α-PD-1 or isotype control immediately prior to immunization with TNP-KLH in alum (Figure 6A). Consistently, sera IgG recovered on day 7 from α-PD-1-treated animals had higher affinity for TNP compared to isotype control-treated animals after immunization (Figure 6B). Little difference in TNP-specific IgG titers were observed between the groups for all subclasses (α-TNP mIgG1, Figure 6C; α-TNP mIgG2a/c, Figure 6D; α-TNP mIgG2b, Figure 6E; α-TNP mIgG3, Figure 6F). Although similar GC B cells was observed (Figure 6G), increased T_FH_ cells (Figure 6H), decreased T_FR_ cells (Figure 6I), and decreased T_FR_: T_FH_ ratio (Figure 6J) were seen following α-PD-1 treatment.

**Figure 6.**
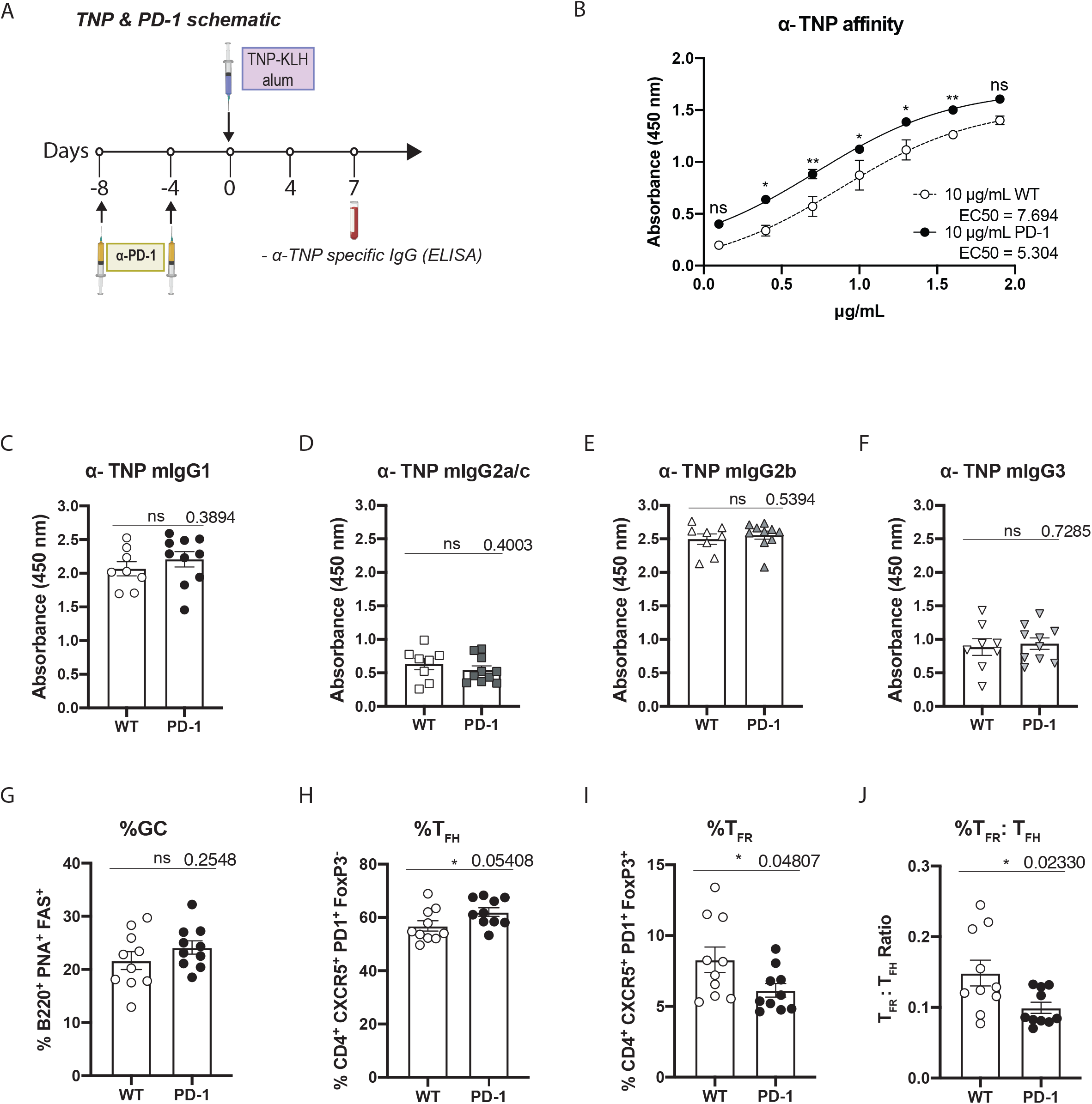
TNP-immunization during PD-1 blockade. (**A)** Schematic of TNP-KLH immunization with α-PD-1 intervention on day −8 and −4. (**B**) TNP-specific affinity constant plot of PBS-treated (white filled) and α-PD-1-treated (black filled) at day 7 after TNP-KLH immunization (*n* = 3-4 mice/group). . Quantification of serum derived TNP-specific mIgG1 (**C**), mIgG2a/c (**D**), mIgG2b (**E**), and mIgG3 (**F**) in PBS-treated (white filled), or α-PD-1-treated (black filled) mice (*n* = 10 mice/group). Flow cytometry-derived percentages of splenocytes in NTN-induction isotype-treated (white circles) and α-PD-1-treated mice (black circles) for germinal center B cells (FAS^+^ PNA^+^) gated on B220^+^ splenocytes (**G**), T follicular helper cells (T_FH_) (CXCR5^+^ PD-1^+^ FoxP3^−^) gated on CD4^+^ cells (**H**), T follicular regulatory cells (T_FR_) (CXCR5^+^ PD-1^+^ FoxP3^+^) gated on CD4^+^ cells (**I**), and T_FR_: T_FH_ ratio (**J**) (*n* = 10 mice/group). Means and standard error of the mean (±SEM) are plotted (*n* = 10 mice/group). Results are representative of at least two independent repeats. *p<0.05, **p<0.01, ***p<0.005, ****p<0.001, ns (not significant), p values are for unpaired *Student’s* t test.

We induced NTN in mice treated with α-PD-1 or isotype control, because this model enabled characterization of sera and tissue-deposited IgG (Figure 7A). α-PD-1 treated animals had worse kidney disease indicated by increased BUN levels on day 7 compared to isotype control treated animals (Figure 7B). Next, the steady-state affinity of kidney-deposited IgG for sheep IgG was examined. IgG recovered from kidneys of α-PD-1 treated animals had higher affinity for sheep IgG compared to control animals (Figure 7C). This trend was true for α-sheep mIgG1 (Figure 7D), α-sheep mIgG2a/c (Figure 7E), and α-sheep mIgG2b (Figure 7F). Importantly, titers of total IgG (mIgG1, Figure 7G; mIgG2a/c, Figure 7H; mIgG2b, Figure 7I; mIgG3, Figure 7J) and sheep IgG-specific IgG (α-sheep mIgG1, Figure 7K; α-sheep mIgG2a/c, Figure 7L; α-sheep mIgG2b, Figure 7M; α-sheep mIgG3, Figure 7N) were unchanged across all IgG subclasses by α-PD-1 treatment. Further, enriched GC B cells (Figure 7O) and T_FH_ cells (Figure 7P), decreased T_FR_ cells (Figure 7Q), and decreased T_FR_: T_FH_ ratio (Figure 7R) (Sommer and Backhed, 2013) were observed following α-PD-1 treatment. Together, abbreviated PD-1 blockade during immunization achieved higher affinity antigen-specific IgG without compromising antigen-specific IgG titers as seen by ABX treatment.

**Figure 7.**
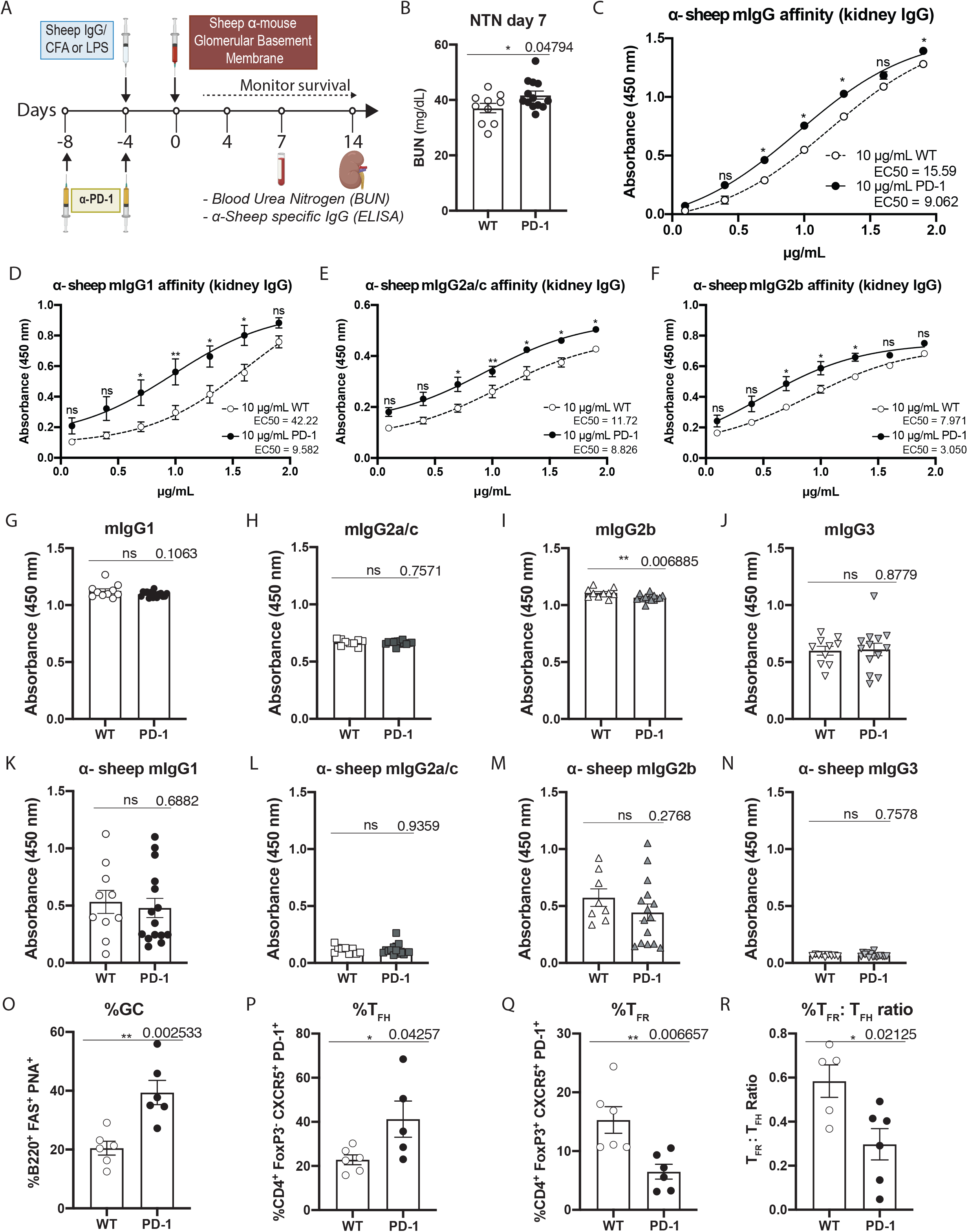
PD-1 blockade during induced nephritis. (**A**) Schematic of NTN-induction with α-PD-1 intervention. (**B**) Blood urea nitrogen (BUN) levels on day 7 of NTN-induction of isotype- (white circles) and α-PD-1-treated mice (black circles) (*n* = 10 mice/group). Sheep IgG-specific affinity plots in NTN-induced isotype- and α-PD-1-treated mice for α-sheep total mIgG (**C**), α-sheep mIgG1 (**D**) (*n* = 3-5 mice/group), α-sheep mIgG2a/c (**E**) (*n* = 3-5 mice/group), α-sheep mIgG2b (**F**) (*n* = 3-5 mice/group). Titers of total mIgG1 **(G)**, mIgG2a/c **(H)**, mIgG2b **(I)**, mIgGg3 **(J)**, and sheep-specific mIgG1 **(K)**, mIgG2a/c **(L)**, mIgG2b **(M)**, and mIgG3 **(N)** 7 days following NTN-induction in isotype-treated (white circles) and α-PD-1-treated mice (black circles) (*n* = 10 mice/group). Flow cytometry-derived percentages of splenocytes in NTN-induction isotype-treated (white circles) and α-PD-1-treated mice (black circles) for germinal center B cells (FAS^+^ PNA^+^) gated on B220^+^ splenocytes (**O**) (*n* = 6 mice/group), T follicular helper cells (T_FH_) (CXCR5^+^ PD-1^+^ FoxP3^−^) gated on CD4^+^ cells (**P**) (*n* = 6 mice/group), T follicular regulatory cells (T_FR_) (CXCR5^+^ PD-1^+^ FoxP3^+^) gated on CD4^+^ cells (**Q**) (*n* = 6 mice/group), and T_FR_: T_FH_ ratio (**R**) (*n* = 6 mice/group). Means and standard error of the mean (±SEM) are plotted. Results are representative of at least two independent repeats. **p<0.05, **p<0.01, ***p<0.005, ****p<0.001, ns (not significant), p values are for unpaired *Student’s* t test.

## Discussion

Productive immune responses coordinate multiple effector arms, often culminating in high affinity IgG. The humoral arm of the immune system is particularly effective in eliciting defense against a number of infections. It is well established that generating high affinity IgG requires T cell help, and more recently the importance of T_FH_ and T_FR_ as critical cells driving the IgG response is appreciated (Wu et al., 2016, Hatzi et al., 2015, Crotty, 2014, Linterman et al., 2011, Crotty, 2011, Zhang et al., 2020, Gowthaman et al., 2019). The balance of these cells ultimately directs antibody production, acting as crucial gatekeepers to IgG responses. Here, we identify two approaches that modulate the balance of T_FH_ and T_FR_, favoring the generation of high affinity IgG.

The microbiota is well established to play a critical role in maintenance of immunological homeostasis and tolerance. Studies have identified metabolites produced by the microbiota, and induction of regulatory T cells as mechanisms through which the immune system is regulated by the microbiota (Takahashi et al., 2020, Maslowski et al., 2009). Our results uncovered a surprising role in this regard, where the microbiota set the balance of cells intimately involved in IgG responses, and specifically IgG affinity, including T_FH_, T_FR_, and GC B cells. Extended ABX leading to transient removal of the microbiota resulted in skewing of these populations in favor of producing high affinity antigen-specific IgG. This held true for two distinct models of immunization, using distinct antigens and adjuvants. The higher affinity IgG resulted in enhanced pathology in the NTN model, despite lower IgG titers and underscores the importance of the quality of the IgG response over quantity.

However, it is unlikely that our extended ABX-regiment be implemented as an immunization strategy for multiple reasons. Indeed, short-term antibiotic regiment was previously shown to impair IgG responses to flu vaccination (Hagan et al., 2019). Therefore, we attempted to functionally recapitulate the effect of extended ABX-treatment, namely increasing T_FH_ cells while decreasing T_FR_ cells. Both of these cells express PD-1 which can be readily targeted by checkpoint inhibition immunotherapy. Indeed, blockade of PD-1 enhanced the T_FR_:T_FH_ ratio in favor of T_FH_ cells. Moreover, this intervention resulted in generation of IgG with higher affinity in two unique immunization models, each with distinct adjuvants. Total and antigen-specific IgG titers were not impaired by α-PD-1 treatment.

Checkpoint inhibition immunotherapy has been a breakthrough for the treatment of many cancers (Gubin et al., 2014, Mamalis et al., 2014). This approach is based on administration of IgG that block inhibitory receptors on immune cells, thereby triggering activating cellular immunity leading to tumor clearance (Wei et al., 2019, Wei et al., 2017, Leach et al., 1996, Iwai et al., 2005, Iwai et al., 2002, Freeman et al., 2000). Although how checkpoint inhibition impacts other aspects of the immune response is in active investigation, reports have indicated intact IgG responses to influenza vaccination in patients undergoing checkpoint therapy, as PD-1 is known to be a negative regulator of B cells (Nishimura et al., 1998)(Ref). We show that PD-1 blockade checkpoint inhibition during immunization with sheep IgG or TNP results in antigen-specific IgG with higher affinity compared to isotype control treatment groups. This intervention did not result in a drop in overall IgG titers, as ABX-treatment did. The availability of PD-1 blockade makes it appealing to speculate it may also improve vaccine responses, enhance monoclonal IgG affinity, or be applied to improve IgG affinity in general. Indeed, potential off-target effects of α-PD-1 are a concern (Nishimura et al., 2001), and a number are associated with cancer immunotherapy. A distinct, and perhaps lower dosing regimen of PD-1 blockade would likely be required. Our experiments showed fewer injections of PD-1 blocking antibodies effectively enhanced the humoral response. Future studies will explore the development of immunological memory responses following PD-1 blockade during vaccination.

## Methods

### *In vivo* Animal Studies

5-6 weeks old C57BL/6 mice for ABX experiments and 7-8 weeks old C57BL/6 and NOD mice were purchased from the Jackson Laboratory and maintained in the animal facility at Massachusetts General Hospital (MGH) under specific pathogen free conditions according to the National Institutes of Health (NIH) guidelines. All animal experiments were approved by and conducted in compliance with the Institutional Animal Care and Use Committee of MGH.

For ABX, 250uL of an antibiotic cocktail consisting of ampicillin (1g/L), metronidazole (1g/L), vancomycin (0.5g/L) and neomycin (1g/L) was given via oral gavage daily as previously described (Rakoff-Nahoum et al., 2004). After daily treatment mice were housed in new cage with autoclaved fresh bedding, food pellets and water. Treatment was initiated at least 3 prior to analysis, immunization, or initiation of inflammation.

To induce nephrotoxic nephritis (NTN), mice were pre-immunized with 200μg of sheep IgG (BioRad) in CFA or LPS via intraperitoneal route, followed by intravenous injection of sheep NTS (Probetex, Inc.) (2μl of serum per gram of mouse) 4 days later. Urea nitrogen (BUN) in sera was measured by the enzyme coupled equilibrium method using a modified urease kit (Stanbio Laboratory). 0.2 mg dose of TNP-KLH immunizations in alum was IP injected and serum was collected 4 and 7 days after. Previous studies have indicated BUN levels for untreated adult mice are 20mg/dL, while NTN-induced animals have BUN levels of 40mg/dL on day 7. Assuming a standard deviation of 9mg/dL for this model, treatment groups of 5/mice would generate p<0.05 at a 95% confidence interval. We therefore randomly assigned five mice to each treatment group for experiments.

KRN TCR transgenic mice on a C57BL/6 background (K/B) were gifts from D. Mathis and C. Benoist (Harvard Medical School, Boston, MA) and were bred to NOD mice to generate K/BxN mice (Korganow et al., 1999). K/BxN serum was prepared as described previously (Kaneko et al., 2006b). Inflammatory arthritis was induced by intravenous injection of K/BxN sera (200μL of pooled K/BxN serum per mouse). Arthritis was scored by clinical examination, and the index of all four paws was added (0 = unaffected, 1 = swelling of one joint, 2 = swelling of more than one joint, 3 = severe swelling of the entire paw) as described (Kaneko et al., 2006b). PBS or ABX treated mice via oral gavage 3-4 weeks prior to K/BxN serum injection.

For PD-1 blockade, 0.2 mg/kg of α-PD-1 (CD279, Biolegend) was administered intraperitoneally. Treatment was initiated 4 days before TNP-KLH in alum immunization. In the nephrotoxic nephritis model, treatment was initiated 8 days and 4 days before pre-immunization with sheep IgG.

### Bacterial stool cultures

Mouse stools samples were weight and diluted 1:10 in sterile PBS based on pellet weight. The stool pellet was then vortexed and spun down in microcentrifuge. 20 μL of mix supernatant was then incubated in LB and BHI plates for 2.5-24 hrs. at 37°C and the CFU/mL were enumerated. For anaerobe incubation, GasPak EZ Gas Generating Pouch Systems (BD) was used as instructed by manufacturer.

### 16S Sequencing

Mouse stool samples were collected, and DNA was extracted as per Powersoil 96 kit (Qiagen). 16S rRNA gene libraries targeting the V4 region of the 16S rRNA gene were prepared by first normalizing template concentrations and determining optimal cycle number by qPCR. To ensure minimal over-amplification, samples were normalized to the lowest concentration sample and were amplified with optimal cycle number for the library construction PCR. Four 25 uL reactions were prepared per sample and each sample was given a unique reverse barcode primer from the Golay primer set (Caporaso et al., 2012, Caporaso et al., 2011a, Caporaso et al., 2011b). Replicates were pooled and cleaned via Agencourt AMPure XP-PCR purification system. Purified libraries were diluted 1:100 and quantified again by qPCR (Two 25uL reactions, 2x iQ SYBR SUPERMix (Bio-Rad, with Read 1 (5’-TATGGTAATTGT GTGYCAGCMGCCGCGGTAA-3’), Read 2 (5’-AGTCAGTCAG CCGGACTACNVGGGTWTCTAAT-3’). Samples were normalized and final pools were sequenced on an Illumina MiSeq 300 using custom index 5’-ATTAGAWACCCBDGTAGTCC GG CTGACTGACT-3’ and custom Read 1 and Read 2 mentioned above. Sequencing data is available from the BioSample database Sequence Read Archive (submission ID: SUB8557868).

### Sequence analysis

Demultiplexed read files were imported into the QIIME2 v2019.7 version of the QIIME software (Bolyen et al., 2019) for analysis. 16S rRNA amplicon primers were first removed with the cutadapt plugin (Martin, 2011). Reads were further denoised, filtered and trimmed with the DADA2 plugin (Callahan et al., 2016). The resulting Amplicon Sequence Variants (ASVs) were classified within QIIME2 using the SILVA v132 reference database (Quast 2013) clustered at 99% sequence identity with a VSEARCH alignment-based method (Bokulich et al., 2018). Prior to downstream analysis, ASVs were filtered to remove counts classified as mitochondria and chloroplasts. Family-level classifications were used for analysis as genus and species-level classifications were incomplete. Stacked bar plots of percent relative abundance of different taxa across samples were generated using the phyloseq package (McMurdie and Holmes, 2013) version 1.30.0. Alpha diversity was measured using Shannon’s Diversity Index (Shannon, 1948) calculated from relative abundance data at the ASV level.

### Total Antibody and IgG subclass ELISA Assays

96-well or 384-well ELISA plates coated with goat α-mouse IgG1, IgG2a/c, IgG2b, IgG3, IgA, IgE and IgM (Bethyl Laboratories) were incubated with 1:1000 diluted sera after blocking with 5% bovine serum albumin. After washing with PBS containing 0.05% Tween 20, the plates were incubated with HRP conjugated anti-mouse IgG-Fc (Bethyl Laboratories), IgA, IgE, IgM (Bethyl Laboratories) and IgG subclasses (Jackson ImmunoResearch). The amount of IgA, IgE, IgG and IgM bound was assessed by 3,3’, 5,5’-tetramethylbensidine (TMB; Biolegend) and the absorbance measured at 450 nm after 2M sulfuric acid addition.

### Antigen-specific IgG ELISA

96-well or 384-well ELISA plates coated with 5μg/mL of sheep IgG or TNP were incubated with 1:500 diluted sera after blocking with 5% bovine serum albumin. After washing with PBS containing 0.05% Tween 20, the plates were incubated with HRP conjugated anti-mouse IgG-Fc (Bethyl Laboratories) and IgG subclasses (Jackson ImmunoResearch). The amount of IgG bound was assessed by 3,3’, 5,5’-tetramethylbensidine (TMB; Biolegend) and the absorbance measured at 450 nm after 2M sulfuric acid addition.

### Preparation of kidney homogenate for IgG purification

Mice were bled on days 4, and 7 after anti-GBM antiserum injection. The serum was separated from the blood by serum gel tubes (BD) and incubated with Protein G high-capacity agarose beads (Thermo Fisher Scientific) for IgG purification. Kidneys were dissected, suspended in 1mL PBS supplemented with protease inhibitor and 2mM EDTA and cut into small pieces before being mechanically homogenized with stainless steel beads and TissueLyser II (Qiagen) for two minutes at 3Hz/s. Homogenate was then diluted 5-fold the volume (PBS with Protein Inhibitor (Thermo) and 2mM EDTA), filtered through 70μm mesh, and centrifuged at 1000×g for 5 min. Supernatant was used to purify IgG with Protein G high-capacity agarose beads.

### Determining affinity of α-sheep IgG and α-TNP IgG

Affinity of antigen-specific IgG was determined as previously described, with minor modifications (Hajighasemi et al., 2004). 96-well or 384-well ELISA plates coated with 20, 10, 5-μg/mL of sheep IgG (Biorad) and purified TNP (BD Pharmingen). Serial dilutions of protein G purified IgG samples were added into coated wells (dilution factor 1:2) ranging from 80-1.25 μg/mL and 50-0.390 μg/mL, respectively. Samples were blocked with 5% bovine serum albumin. After washing with PBS containing 0.05% Tween 20, the plates were incubated with HRP conjugated anti-mouse IgG-Fc (Bethyl Laboratories) and IgG subclasses (Jackson ImmunoResearch). Bound IgG was assessed by 3,3’, 5,5’-tetramethylbensidine (TMB; Biolegend) and the absorbance measured at 450 nm after 2M sulfuric acid addition. The amount of antibody bound to antigen was represented as a sigmoidal curve of absorbance vs the logarithm of the antibody concentration per well (Hajighasemi et al., 2004). The antibody concentration resulting in 50% of maximum absorbance value at a particular antigen coating concentration was selected for the affinity calculation via Prism and reported after analysis as ‘EC50’.

### Flow Cytometry

Cells were collected for flow cytometry by filtering spleen through a 40 μm nylon membrane. Surface staining: CD4, CD8, CD3, CD21/35, B220, CD19, FAS, IgM, IgD, CD23, CXCR5, PD-1, PNA, CD11b, CD11c, GL7 and GR1from BD Pharmingen and Biolegend. Intracellular staining was performed using FoxP3 Fix/Perm buffer set (Biolegend) per manufacturer’s instructions. Cells were run on an LSRII (BD Biosciences) and CytoFLEX S (Beckman Coulter). Analysis was performed with FlowJo (BD Biosciences).

### Quantification and Statistical Analysis

Data from immunization experiments was analyzed in GraphPad Prism: *p<0.05, **p<0.01, ***p<0.005, ****p<0.001 as determined by *Student’s t* test or by two-way ANOVA followed by Tukey’s posthoc.

## Acknowledgements

The authors would like to thank Kai-Ting Shade, Maya Kitaoka, Kate Jeffrey and Jim Moon for careful reading and constructive comments of this manuscript, and the Broad Institute’s Microbial ‘Omics Core and Genomics Platform for sample processing and sequencing data generation. This work was supported by NIH awards R01AI153441 and DP2AR068272 to RMA and P30 DK043351 to RX. JP performed the experiments, collected samples, and analyzed data. HK, AG analyzed 16S sequencing data, RX and RMA conceived of the experiments, and JP and RMA wrote the manuscript.

## Supplemental Data

Key Resources Table.

Key Resources Table
| ANTIBODIES | SOURCE | IDENTIFIER |
| --- | --- | --- |
| APC-Cy7 anti-mouse CD4 | Biolegend | Cat#: 100413 |
| PE-Cyanine 5.5 anti-mouse CD45 | eBioscience | Cat#: 35-0451 |
| PE-Cy7 anti-mouse GL7 | Biolegend | Cat#: 144620 |
| Alexa Flour 647 anti-mouse CD279 (PD-1) | Biolegend | Cat#: 135230 |
| Alexa Flour 488 anti-mouse FoxP3 | Biolegend | Cat#: 320012 |
| PE anti-mouse CXCR5 | Biolegend | Cat#: 145503 |
| PE hamster anti-mouse CD95 (Fas) | BD Pharmingen | Cat#: 554258 |
| Lectin PNA, Alexa Flour 488 | Molecular Probes | Cat#: L-21409 |
| APC anti-mouse IgM | Biolegend | Cat#: 406509 |
| PE-Cy5.5 anti-mouse CD45 | Biolegend | Cat#: 35-0451-80 |
| PerCP anti-mouse IgD | Biolegend | Cat#: 405709 |
| FITC anti-mouse CD21/35 | Biolegend | Cat#: 123407 |
| Sheep IgG | BioRad | Cat#: PSP01 |
| PE anti-mouse CD23 | Biolegend | Cat#: 101607 |
| Biotin anti-mouse B220 | Biolegend | Cat#: 103204 |
| Cy5 Streptavidin | Biolegend | Cat#: 405209 |
| Trustain FcX (anti-mouse CD16/32) | Biolegend | Cat#: 101320 |
| PerCP-Cyanine 5.5 anti-mouse CD8a | eBioscience | Cat#: 45-0081 |
| GoInVivo Purified anti-mouse CD279 (PD-1) antibody | Biolegend | Cat#: 135234 |
| Purified mouse anti-TNP | BD Biosciences | Cat#: 554054 |
| Biotin-SP-conjugated AffiniPure anti-mouse IgG, Fcγ subclass 1 specific | Jackson ImmunoResearch | Cat#: 115-065-205 |
| Biotin-SP-conjugated AffiniPure anti-mouse IgG, Fcγ subclass 2b specific | Jackson ImmunoResearch | Cat#: 115-065-207 |
| Biotin-SP-conjugated AffiniPure anti-mouse IgG, Fcγ subclass 2c specific | Jackson ImmunoResearch | Cat#: 115-065-208 |
| Biotin-SP-conjugated AffiniPure anti-mouse IgG, Fcγ subclass 3 specific | Jackson ImmunoResearch | Cat#: 115-065-209 |
| Goat anti-Mouse IgA Antibody Affinity Purified | Bethyl Laboratories, Inc. | Cat#: A90-103A |
| Goat anti-Mouse IgA Antibody HRP Conjugated | Bethyl Laboratories, Inc. | Cat#: A90-103P |
| Goat anti-Mouse IgE Antibody Affinity Purified | Bethyl Laboratories, Inc. | Cat#: A90-115A |
| Goat anti-Mouse IgE Antibody HRP Conjugated | Bethyl Laboratories, Inc. | Cat#: A90-115P |
| Goat anti-Mouse IgM cross-adsorbed Antibody Affinity Purified | Bethyl Laboratories, Inc. | Cat#: A90-201A |
| Goat anti-Mouse IgM cross-adsorbed Antibody HRP Conjugated | Bethyl Laboratories, Inc. | Cat#: A90-201P |
| Goat anti-Mouse IgG-Fc Fragment cross-adsorbed Antibody HRP Conjugated | Bethyl Laboratories, Inc. | Cat#: A90-231P |
| FITC anti-mouse CD11b | Biolegend | Cat#: 101205 |
| FITC anti-mouse CD11c | Biolegend | Cat#: 117305 |
| FITC anti-mouse GR1 | Biolegend | Cat#: 108405 |
| FITC anti-mouse CD3. | Biolegend | Cat#: 100203 |
| **CHEMICALS, PEPTIDS, RECOMBINANT PROTEINS** | **SOURCE** | **IDENTIFIER** |
| Ampicillin sodium salt | RPI (Research Products International) | Cat#: A40040-10.0 |
| Metronidazole | Sigma-Aldrich | Cat#: M3761-25G |
| Neomycin trisulfate salt | Sigma | Cat#: 6386-25G |
| Vancomycin Hydrochloride | Alvogen | Cat#: 47781-597-07 |
| Pierce™ Protein G Plus Agarose | Thermo Fisher | Cat#: 22852B |
| NTS (Sheep Anti-Rat Glomeruli (Anti-GBM) Serum) | Probetex | Cat#: PTX001-S |
| TMB Substrate | Biolegend | Cat#: 421101 |
| Protease Inhibitor Mini Tablets | Thermo Fisher | Cat#: 88665 |
| FoxP3 Fix/Perm Buffer Set | Biolegend | Cat#: 421403 |
| Lipopolysaccharide from *E.coli* 055:B5 | MD BioProducts | Cat#: MDLPS<br>Lot No: 20317A2 |
| TNP-KLH (Keyhole Limpet Hemocyanin) | BioSearch Technologies | Cat#: T-5060 |
| Complete Freund's Adjuvant | BD | Cat#: 263810<br>Lot No: 6231897 |
| Alum-Imject Adjuvant | Thermo Scientific | Cat#: 77161<br>Lot No: TC263121 |
| PowerSoil, DNA Isolation Kit | Qiagen | Cat#: 12888-50 |
| GasPak EZ Anaerobe Gas Generating Pouch System | BD | Cat#: 260683 |
| Luria agar | RPI (Research Products International) | Cat#: L24025-1000 |
| Brain Heart Infusion Broth | Fluka Analytical | Cat#: 53286-500G |
| HRP-Avidin | Biolegend | Cat#: 405103 |
| **ASSAY KITS** | **SOURCE** | **IDENTIFIER** |
| Stanbio Urea Nitrogen (BUN) Liqui-UV Test | Stanbio Laboratory | Cat#: 2020-0430<br>Lot No: 171161 |
| **EXPERIMENTAL MODELS: Organisms/Strains** | **SOURCE** | **IDENTIFIER** |
| Mouse: C57BL/6J | The Jackson Laboratory | RRID:IMSR_JAX:000664 |
| **SOFTWARE AND ALGORYTHMS** | **SOURCE** | **LINK** |
| GraphPad Prism 7 | GraphPad Software, Inc. | https://www.graphpad.com/scientific-software/prism/ |
| QIIME2 | QIIME2 | https://qiime2.org/ |

---

**Figure S1.**
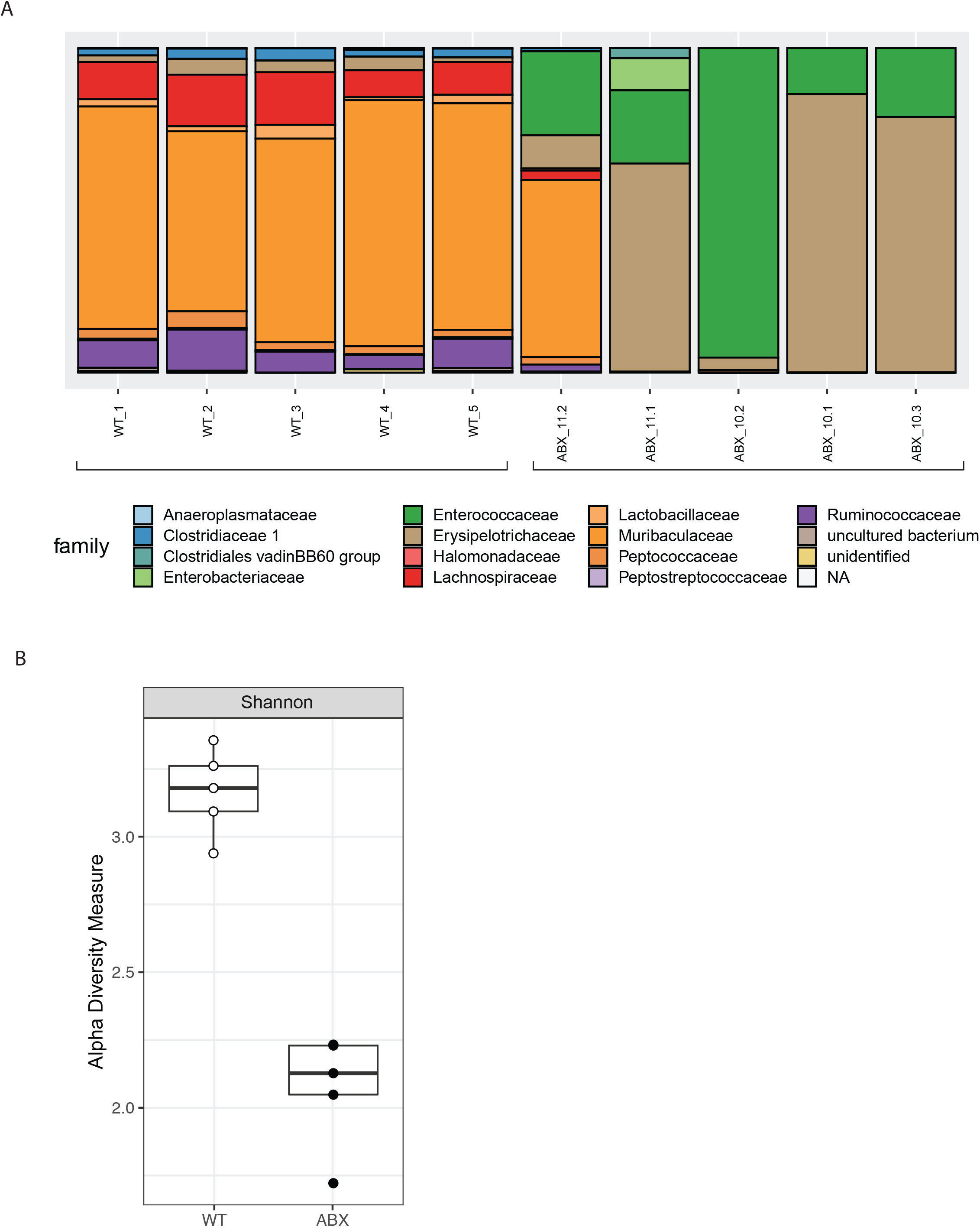
Analysis of 16S Sequencing of microbiota. **(A)** Relative percentage of fecal microbial families identified by 16S sequencing. **(B)** Diversity of fecal microbial populations in WT control and ABX mice as determined by 16S sequencing.

**Figure S2.**
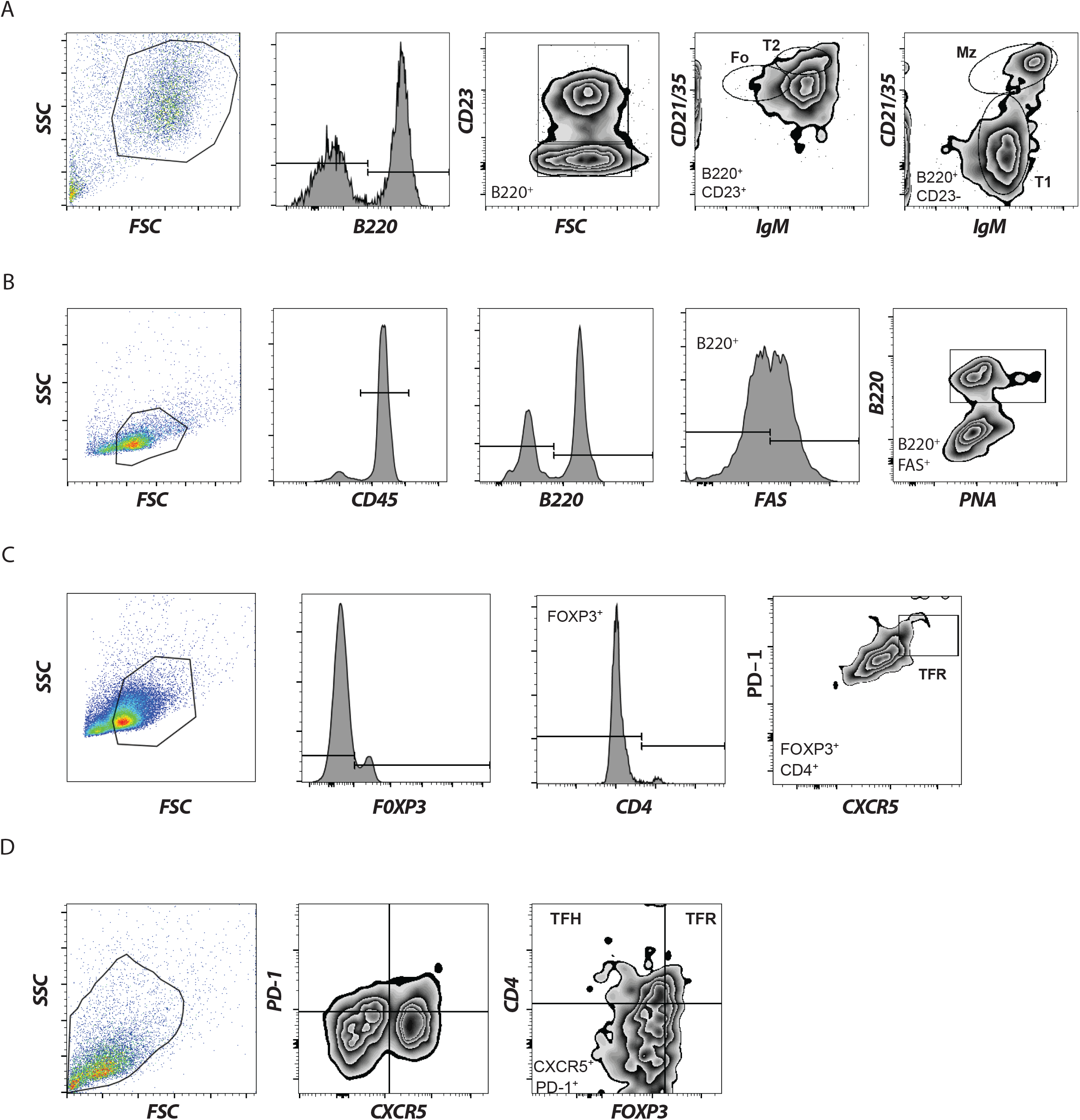
Flow cytometric gating strategies for lymphocyte populations. **(A)** To identify B cell populations, forward scatter (FSC) and side scatter (SSC) lymphocytes were first gated, by B220^+^ cells, then CD23^+^ cells, then CD21/35 and IgM were used to identify Follicular B cells (Fo, B220^+^ CD23^+^ CD21/35^+^, IgM^−^), Transition 2 cell (T2, B220^+^ CD23^+^ CD21/35^+^, IgM^+^), Transition 1 cells (T1, B220^+^ CD23^−^ CD21/35^−^, IgM^+^), and Marginal Zone B cells (Mz, B220^+^ CD23^−^ CD21/35^+^, IgM^+^). **(B)** To identify Germinal Center B cells, forward scatter (FSC) and side scatter (SSC) lymphocytes were gated, followed by CD45+ cells, followed by B220^+^ PNA^+^. **(C)** To identify T follicular regulatory cells (CD4^+^ FoxP3^+^ PD-1^+^ CXCR5^+^), forward scatter (FSC) and side scatter (SSC) lymphocytes were gated, followed by FoxP3, CD4, PD-1 and CXCR5. (D) T_FR_ and T_FH_ ratios were determined by forward scatter (FSC) and side scatter (SSC) lymphocyte gating, followed by PD-1^+^ CXCR5^+^ CD4^+^ cells that expressed or did not expressed FoxP3 (T_FR_ or T_FH_, respectively.

**Figure S3.**
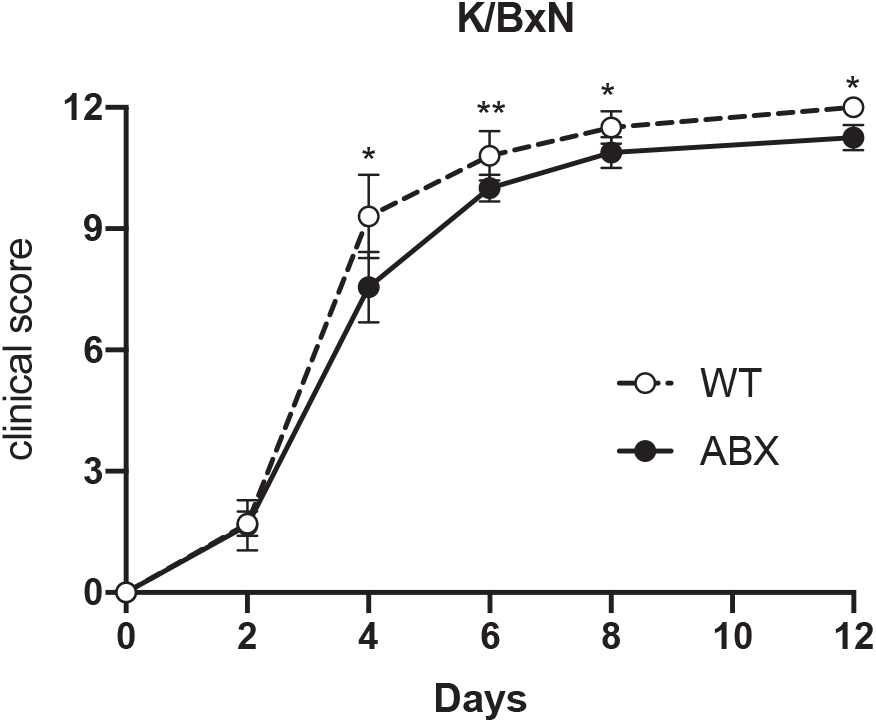
Passive K/BxN sera transfer to WT or ABX mice. Clinical scores of WT or ABX mice administered K/BxN sera over several days. Means and standard deviations are shown. *p<0.05, **p<0.01 as determined by ANOVA followed by a Tukey’s posthoc test.

**Figure S4. Characterization of the α-sheep IgG response for NTN-passive transfer.** Sheep IgG specific titers for mIgG1 (A), mIgG2a/c (B), mIgG2b (C), and mIgG3 (D) 7 days after NTN induction in WT controls (white) and ABX (black) mice. E. Schematic of recovery and characterization of kidney-deposited IgG. **(F)** Anti-sheep IgG1 affinity plot of total IgG antibodies recovered after protein G purification from serum at day 9. **(G)** Anti-sheep IgG2a/c affinity plot of total IgG antibodies recovered after protein G purification from serum at day 9. **(H)** Anti-sheep IgG2b affinity plot of total IgG antibodies recovered after protein G purification from serum at day 9. Means and standard deviation are plotted. Results are representative of at least two independent repeats. *p<0.05, **p<0.01, ***p<0.005, ****p<0.001, ns (not significant), p values are for unpaired *Student’s* t test.

